# Left Out: The Effect of Handedness on fMRI Activation in Memory Paradigms

**DOI:** 10.1101/2023.08.17.553587

**Authors:** Loriana Goulding, Isaac Hamm, C. Brock Kirwan

**Affiliations:** Department of Psychology, Brigham Young University; MindCORE Neuroimaging Facility, University of Pennsylvania

**Author notes:** Correspondence should be addressed to: Brock Kirwan, 3401 Grays Ferry Ave., Building 250 Ste 140, Philadelphia, PA 19146.

**Keywords:** memory, handedness, lateralization, fMRI, hippocampus

## Abstract

About 11% of the population is left-handed, a significant minority of the potential participant pool for functional MRI (fMRI) studies. However, convention in fMRI research dictates these potential participants be excluded due to the supposition that left-handed (LH) people may have different lateralization of neural functioning than right-handed (RH) people. This difference in lateralization may cause different areas of the brain to be activated by the same task. The current study investigates the lateralization differences between LH (n=54) and RH (n=55) participants during encoding and recognition memory tasks for verbal and non-verbal stimuli. Additionally, we measured participants’ language laterality index by administering a semantic fluency task. We found no difference between groups’ behavioral memory performance for either verbal or non-verbal stimuli. Similarly, we found no group differences in fMRI activation for either verbal or non-verbal material at encoding or retrieval. When we split groups according to language lateralization, we observed more activation in right-lateralized participants for word retrieval only. To measure whether including LH in fMRI studies would dilute results, we performed Monte Carlo simulations with subsets of our data for sample sizes from n=10 to 50 with the number of LH in each sample ranging from k=0 to 9. Similarity metrics between activation maps for the all-RH sample and samples that include LH participants were robust, especially for large sample sizes (e.g., Dice similarity coefficients >0.9 when n=50). Taken together, these results indicate that the inclusion of left-handed participants does not have a strong detrimental effect on memory-related fMRI activation. On this basis we advocate for the inclusion of left-handed participants in cognitive neuroscience of memory research.

## Introduction

Traditionally, left-handed (LH) individuals have been excluded from fMRI research based on the idea that asymmetries in brain function, or brain lateralization, differs between LH and right-handed (RH) people to the degree that including them in a sample would significantly interfere with the results. Although LH individuals make up approximately 11% of the population (de Kovel et al., 2019; Gilbert & Wysocki, 1992), only 18% of fMRI studies report including one or more LH participants, representing only 3-4% of studied participants (Bailey et al., 2020). As fMRI analyses are largely comparative by nature, including subjects with differing underlying functional organization patterns may skew results by introducing variability in activation beyond that which is caused by the phenomena of interest. However, excluding LH individuals may lead to systematic bias in studies where handedness is not significantly related to levels of activation in regions of interest.

Lateralization of brain function has been observed going back to classic lesion studies from the 19^th^ century where clinicians observed that patients with lesions to the left hemisphere resulted in language impairments, leading to the conclusion that language function is localized to the left hemisphere in almost all RH people (Damasio & Geschwind, 1984). In some areas of fMRI study, the brain lateralization for LH and RH individuals has been shown to be especially divergent, which often leads to LH individuals being excluded from these types of research (e.g., language and motor-skills based research; Bailey et al., 2020; McManus, 2019). One area of the literature that lacks substantial research is the brain lateralization of memory processes in LH and RH people. Exclusion from fMRI studies based on handedness exacerbates the already difficult participant selection process, as it excludes more than one-tenth of the US population (Gilbert & Wysocki, 1992). Moreover, given that LH individuals make up a sizable portion of the population, the blanket exclusion of LH individuals from fMRI research may limit the generalizability of the research to the general population and is ethically questionable (Bailey et al., 2020; Willems et al., 2014).

Although certain areas of the brain have been shown to have differences in structural lateralization between LH and RH individuals—for example, the fusiform face area, intraparietal sulcus, and right ventral and dorsal premotor cortex—many areas of the brain have not shown this handedness-driven lateralization for either structural or functional measures (Frässle et al., 2016; Martin et al., 2011). In fMRI language research, LH individuals are typically excluded because research has concluded that they are more likely than RH individuals to exhibit more language-related activation in the right than left hemisphere, commonly referred to as “atypical language lateralization” (Szaflarski et al., 2002, but see Szaflarski et al., 2012). However, brain lateralization is not as straightforward as it may first appear. In facial recognition research, for example, early face processing as well as creation and comparison of facial representations have been shown to be more efficient in the right hemisphere (Rhodes, 1985), and left visual field superiority has been shown for facial recognition (Hilliard, 1973). Later research also demonstrated that both hemispheres participate differentially in facial recognition (Badzakova-Trajkov et al., 2010; Meng et al., 2012). Given recent data indicating that the hippocampus preferentially responds to information in the contralateral visual field (Silson et al., 2021) — consistent with the theory of lateralized functioning in the hippocampus—there is a basis for testing whether handedness would affect activation in memory research. Furthermore, several models of episodic memory are explicitly lateralized, e.g., the Hemispheric Asymmetries of Memory (HERA) model (Habib et al., 2003). Nevertheless, the effects of handedness on the lateralization of memory functions have yet to be thoroughly explored.

Golby et al. (2001) concluded that the most important aspect of memory-encoding lateralization was related to the verbalizability of the stimuli by observing that verbal encoding resulted in left-lateralized activation in the inferior prefrontal cortex and in the medial temporal lobe, whereas visual pattern encoding activated the right inferior prefrontal cortex and the right medial temporal lobe, and scenes and faces resulted in fairly similar activation in both regions. However, Golby et al. (2001) did not use handedness as a variable in their research and rather examined lateralization differences of memory functions in an all-RH sample. While Cuzzocreo et al. (2009) found a difference in activation of lateralization in fMRI memory research on the basis of handedness, their study only tested verbal memory in participants above the age of 50, thus leaving open the question of whether memory for non-verbal stimuli would be affected and whether the results generalize to other age groups. Additionally, while Cuzzocreo et al. attribute the difference found in activation to differences in lateralization based on handedness, no language laterality index (LI_lang_) was calculated for participants to rule out other explanations. Because verbal and spatial memory functions have been shown to change in lateralization due to aging (Reuter-Lorenz et al., 2000), age could be an important factor in lateralization.

Our study builds on previous research by including a younger population, measures of both verbal and nonverbal memory, and LI_lang_ calculated for each participant. Consistent with previous research, we hypothesized that when performing verbal and nonverbal memory tasks, there would be lateralized activation differences in memory-dependent contrasts between LH and RH groups at both encoding and retrieval. Given the large body of research showing handedness-related lateralization of language, and the mixed results regarding lateralization in facial processing related to handedness (Willems et al., 2014), we also hypothesized that at both encoding and retrieval (tested separately) there would be differential memory activation between groups such that the lateralized activation differences would be stronger for words than for faces.

Our second research question examined whether including LH individuals would lessen any activation effects observed with an all-RH sample (i.e., does adding LH individuals to an analysis that is significant for RH individuals make the effect fail to reach significance?) and if so, how many LH participants could be added into an analysis before the effect becomes diluted enough to fail to be detected. We hypothesized that at both encoding and retrieval we would see the difference in activation for memory effects decrease (fail to reach significance) as more RH participants were replaced by an LH counterpart in the analysis.

## Methods

### Participants

One-hundred and twenty participants (61 LH and 59 RH individuals) were recruited from the Brigham Young University campus and surrounding community in Provo, UT, USA. Inclusion criteria required participants to be in good overall health and to have no history of physiological or neurological disorders. Participants were prescreened for handedness using the 10-item Edinburgh Handedness Inventory (EHI), which asks participants to self-report their preferred hand for a variety of manual activities. Participants responded on a 5-point Likert scale with options “always left”, “mostly left”, “no preference”, “mostly right”, and “always right”, which were scored as -10, -5, 0, 5, and 10, respectively. Although recent research suggests poor correspondence between the EHI and other measures of handedness (Ruck & Schoenemann, 2021), it is nevertheless the most widely used instrument for assessing handedness in the neuroimaging literature (McManus, 2019; Yeung & Wong, 2022). Eligibility for inclusion required composite EHI scores of ≤ -40 for the LH and ≥ 40 for the RH group to ensure a clear handedness orientation (Edlin et al., 2015; Oldfield, 1971). Safety screening was also conducted to ensure participants were eligible for MRI scanning. Participants received either $20 or a ¼-scale 3D-printed model of their brain as remuneration for their participation. The Brigham Young University Institutional Review Board approved all study protocols and procedures, and participants gave informed consent before participating. Two participants were excluded from both behavioral and neuroimaging analyses due to failure to follow instructions; two were excluded due to poor memory performance (d’ scores more than 2 *SD*s below the mean); one was excluded due to equipment failure; two were excluded due to poor MRI image quality; and one was excluded for an incidental neurological finding. Three participants had excessive motion during MRI scanning (defined as more than 20% of TRs with movement per scan run) and were excluded from all runs; three additional participants were excluded from the word test condition; and six were excluded from the face test condition. Accordingly, our final sample size was N=109 (LH=54, RH=55; age range 18-47, mean 21.8, SD 3.4; 61 female).

### Procedure

Participants completed, in order, a word encoding task, a face encoding task, a semantic fluency task (Brumer et al., 2020), a word retrieval task, and a face retrieval task while undergoing fMRI scanning (see Figure 1). In the encoding tasks, participants were shown 100 faces and then 100 words in separate blocks and were asked to determine whether they considered each item presented pleasant or unpleasant (via an MR-compatible response device) to promote deeper stimulus encoding (Guerin & Miller, 2009). Each stimulus was presented for 2.5 seconds followed by a fixation cross for 0.5-1.5 seconds (jittered). Stimulus order was randomized within each stimulus type. Faces were selected from a database made to be broadly representative of adult age, gender, and race (Minear & Park, 2004). Words were nouns selected from the MRC Psycholinguistics Database (Coltheart, 1981) with high concreteness (≥600) and familiarity (≥400) ratings.

**Figure 1.**
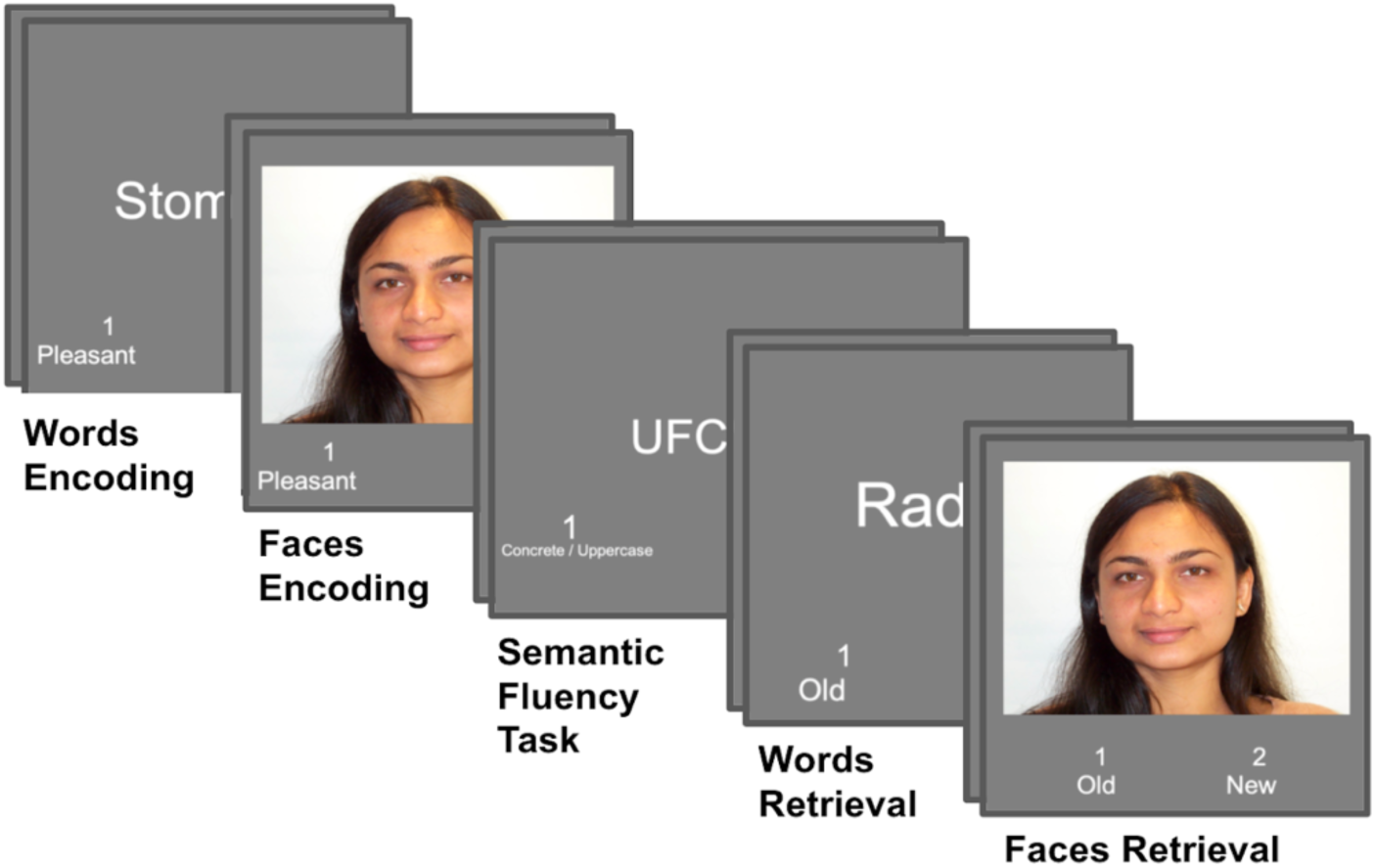
Participants completed encoding, semantic fluency, and retrieval tasks while undergoing fMRI scanning. In each task, each stimulus appeared for 2,500ms with an interstimulus interval jitter of 500-1,500ms, during which a single fixation cross was presented in the center of the screen. Task order was the same for all participants, but stimuli were randomized within each task across participants.

Next, we administered a semantic fluency task that has been shown to demonstrate activation in frontal and temporoparietal regions (Bradshaw, Thompson, et al., 2017), which we used to calculate a LI_lang_ for each participant. The task consisted of six 30-second cycles alternating between activation and baseline conditions (Brumer et al., 2020; Harrington et al., 2006). For the activation condition, participants were shown abstract and concrete words in a random order and were asked to identify whether each word was abstract or concrete.

Participants were given the definition and examples of “abstract” vs “concrete” words prior to scanning. Words were again selected from the MRC Psycholinguistics Database (Coltheart, 1981). All stimuli had a familiarity rating ≥ 400; abstract words had concreteness ratings ≤ 400; and concrete words had concreteness ratings ≥ 600. For the baseline condition, participants were shown randomized strings of letters in all upper- or lowercase and were asked to identify whether the strings were upper- or lowercase.

Finally, participants performed memory retrieval tasks for faces then words. In both tasks, participants were shown 100 target stimuli and 100 novel foil stimuli. The order of the individual words and faces was again randomized. Participants were asked to identify the previously seen (“old”) and novel (“new”) stimuli by pressing one of two buttons. Each stimulus was presented for 2.5 seconds followed by a fixation cross for 0.5-1.5 seconds (jittered). The retrieval tasks were each broken up into two scan runs of 100 trials each.

Prior to being positioned in the MRI scanner, participants were given instructions regarding the behavioral tasks they would be performing and information concerning MRI safety measures. Participants used a four-button response device (Current Designs Inc.; Philadelphia, PA) and made button presses with the pointer finger and middle finger of the right hand. Stimuli were displayed on an MRI compatible LCD monitor (BOLDscreen; Cambridge Research Systems; Rochester, UK) located at the head-end of the MRI scanner and viewed via a mirror mounted to the head coil. Instructions regarding which button corresponded with which stimulus type were listed on each slide throughout each phase of the study.

### Imaging Procedures

A Siemens 3 Tesla TIM Trio scanner (Erlangen, Germany) with a 32-channel head coil was used for all MR scanning. We collected a T1-weighted structural scan in addition to echo-planar imaging (EPI) scans for each of the tasks. T1-weighted MP-RAGE parameters: TR = 1900 ms; TE = 4.92 ms; flip angle = 9°; field of view = 256 mm; slice thickness = 1 mm; 176 slices; voxel resolution = .97 × .97 × 1.0 mm; 1 average. EPI scans used a T2*-weighted pulse sequence utilizing a multiband (MB) technique (Xu et al., 2013) with the following parameters: 72 interleaved slices; TR = 1800 ms; TE = 42 ms; flip angle = 90°; field of view = 180 mm; slice thickness = 1.8 mm; voxel resolution = 1.8 × 1.8 × 1.8 mm; MB factor = 4. Each functional run had 195 acquisitions except for the semantic fluency task, which had 200. The first four TRs of each run were discarded to allow for T1 equilibration. All (anonymized) MRI scan data are available at https://openneuro.org/datasets/ds004589.

### Data Analysis

#### Pre-Processing

Structural and functional MRI data were converted from DICOM to NIfTI format using program *dcm2niix* (Li et al., 2016) and structural scans were de-faced for anonymity. Data were preprocessed using AFNI program *afni_proc.py* (Taylor et al., 2024). Briefly, fMRI data were distortion corrected, slice-time corrected, co-registered to the structural scan, normalized to the MNI atlas, motion corrected, and masked. Functional data were blurred with a 4mm FWHM Gaussian blur and scaled by the mean of the overall signal for each run. Functional volumes with large motion events, defined as TRs with a Euclidean norm (ENORM) of the temporal derivative of motion estimates (rotations and translations) greater than 0.3, along with TRs immediately before a motion-contaminated TR, were excluded from single-participant regression analyses.

We created separate single-subject regression models for each phase of the experiment. Behavioral regressors for both encoding tasks coded for subsequent hits, subsequent misses, and trials of no interest (e.g., non-responses at either encoding or retrieval). Behavioral regressors for both retrieval tasks coded for hits, misses, correct rejections (CRs), false alarms (FAs), and trials of no interest. Events in all encoding and retrieval tasks were modeled as a canonical hemodynamic response function convolved with a boxcar function of 2.5-second duration. The semantic fluency task was modeled as a block-design with a 30-second boxcar for the active task condition convolved with the canonical hemodynamic response function. All single-subject regression models included six regressors for motion (three translation, three rotation) and polynomial regressors coding for run (in the case of the retrieval tasks that had two scan runs each) and scanner drift. To identify memory-related activation, we performed contrasts for subsequent hits vs. subsequent misses for the encoding data and for hits vs. correct rejections for the retrieval data. The resulting statistical maps were then entered into group analyses as described below.

#### Language Laterality Index and Group Analyses

To define the LI_lang_, we first conducted a meta-analysis of fMRI papers using NeuroSynth (https://neurosynth.org) for the term “language”. The association test z-map, which identifies regions that are preferentially active in studies that load heavily on the term “language” (see Yarkoni et al., 2011), was resampled to match our functional resolution, thresholded, and then left-right mirrored to be symmetrical across the midline (Wilke & Lidzba, 2007). Masks are available at https://osf.io/f568s/overview?view_only=4f53cd74ad5646e68272ee9ff9e6b4a4.

Consistent with previous studies (Bradshaw, Bishop, et al., 2017; Brumer et al., 2020; Harrington et al., 2006; Jensen-Kondering et al., 2012), we focused on the cluster of voxels identified in the left and right temporal lobes (primarily superior temporal gyrus) to calculate the LI_lang_. The LI_lang_ was calculated for each subject by z-transforming the single-subject regression results for the semantic fluency task and then thresholding the output at p < .05 (Bradshaw, Thompson, et al., 2017). We then extracted the mean activation for all voxels within the left and right temporal lobe language clusters that were more active for the task than the baseline conditions. The LI_lang_ was calculated as (L-R)/(L+R), where L is the mean activation in the left temporal lobe and R is the mean activation in the right temporal lobe.

Whole-brain analyses were performed on the encoding and retrieval data using independent-samples t-tests (AFNI program *3dttest++*) as described below. All analyses were confined to voxels identified as within signal coverage for 80% of individual subjects. Multiple comparisons were corrected using permutation testing based on residuals at the group level (Cox et al., 2017) with voxel-wise p<.01 and spatial extent threshold k>400 for FWE p<.05. Unless otherwise noted, all t-test comparisons reported are two-tailed.

We also conducted anatomical ROI analyses focused on the medial temporal lobe (MTL). Regions of interest (ROI) were defined manually by an expert rater (CBK) for the hippocampus, entorhinal cortex, perirhinal cortex, and parahippocampal cortex on the MNI template (Insausti et al., 1998) and resampled to resolution of the functional scans. Anatomical masks were limited to voxels with 80% signal coverage as in the whole-brain analyses. Mean beta contrasts were extracted for each anatomical ROI and each task and were then investigated with 2 (group) × 2 (hemisphere) × 4 (ROI) ANOVAs. Where appropriate, Greenhouse-Geisser corrected degrees of freedom are reported.

#### Monte Carlo Simulations

To test the hypothesis that including LH participants would increase the variability of results compared to all-RH samples, we performed Monte Carlo simulations where we randomly selected subsets of participants with sample sizes between 10 and 50 (i.e., n=10, 15, …, 50) and included between 0 and 9 (i.e., k=0, 1, …, 9) LH participants. We calculated 100 activation maps and then took the average t-statistics across all iterations for each (n,k) combination and for each task condition (for a total of 9,000 iterations for each task condition). We thresholded average t-maps corresponding to p<.05 (one-tailed) and then calculated Dice similarity coefficients (DSC) between the activation maps with k=0 LH participants (i.e., the all-RH activation maps) and all values of k for each sample size.

Finally, to further examine the effect of adding participants with “atypical lateralization” (i.e., negative LI_lang_ scores) to the Monte Carlo simulation samples, we calculated the effect of mean LI_lang_ on DSC for each pairwise comparison of all unique draws (excluding self-comparisons) in the k=0 and k=5 conditions for the largest sample size (n=50). Since the Monte Carlo simulations involved random sampling with replacement, there was varying overlap between random draws and more overlap in the k=0 condition than the k=5 condition.

Accordingly, for each pair of draws, we calculated both subject overlap, quantified as the Jaccard index (the number of subjects common to both draws divided by the number of subjects in either draw), and the mean difference in LI_lang_ between participants composing each draw. We fit a linear mixed-effects model predicting pairwise DSC as a function of subject overlap (Jaccard index), LI_lang_ difference between draws, inclusion of LH participants (k=0 or k=5), and their interaction, with task included as a fixed effect to account for baseline differences in map similarity across task conditions: DSC ∼ Jaccard + ΔLI × k + task + (1 | draw_A) + (1 | draw_B). Crossed random intercepts for each unique draw (identified by its role as the lower- or higher-indexed member of a canonicalized unordered pair) accounted for the non-independence of DSC values sharing a common draw. Models were fit via restricted maximum likelihood using the *lme4* package (Bates et al., 2015) in R.

Analysis scripts and masks are available at https://osf.io/f568s/overview?view_only=4f53cd74ad5646e68272ee9ff9e6b4a4.

## Results

### Behavioral Performance

Proportions of responses in the words and faces recognition tasks along with discriminability (d’) scores are reported in Table 1. To examine whether there were group differences in memory performance, we performed a linear mixed-effects regression on d’ scores with stimulus type and handedness as fixed effects. There was no main effect of group (F(1,107) = 3.392, p = 0.068) and no group × stimulus type interaction (F(1,107) = 0.130, p = 0.719). There was a significant main effect of stimulus type (F(1,107) = 408.331, p < 0.001) driven by much higher memory performance for words than for faces.

**Table 1.** Mean (SD) proportions of responses and d’ scores.

|  | Words |  |  | Faces |  |  |
| --- | --- | --- | --- | --- | --- | --- |
| | Hits | CRs | $d'$ | Hits | CRs | $d'$ |
| RH | .92 (.06) | .89 (.08) | 2.86 (.74) | .70 (.13) | .80 (.12) | 1.48 (.43) |
| LH | .91 (.06) | .88 (.07) | 2.67 (.59) | .65 (.14) | .80 (.12) | 1.34 (.55) |

*Note.* RH=right-handed; LH=left-handed; CRs=correct rejections; $d'$ =discriminability score calculated as the $z(\text{hits})-z(\text{false alarms})$ .

### Neuroimaging Results

#### Language Laterality Indices

There was a significant difference between LH and RH participants in LI_lang_ (LH = 0.527 (0.547); RH = 0.764 (0.272); t(107) = 2.865, p = 0.005, 95%CI [-0.400, -0.073]). All the RH participants were left-lateralized (i.e., had positive LI_lang_ values), and most of the LH (45 out of 54 or 83.3%) were also left-lateralized (see Figure 2).

**Figure 2.**
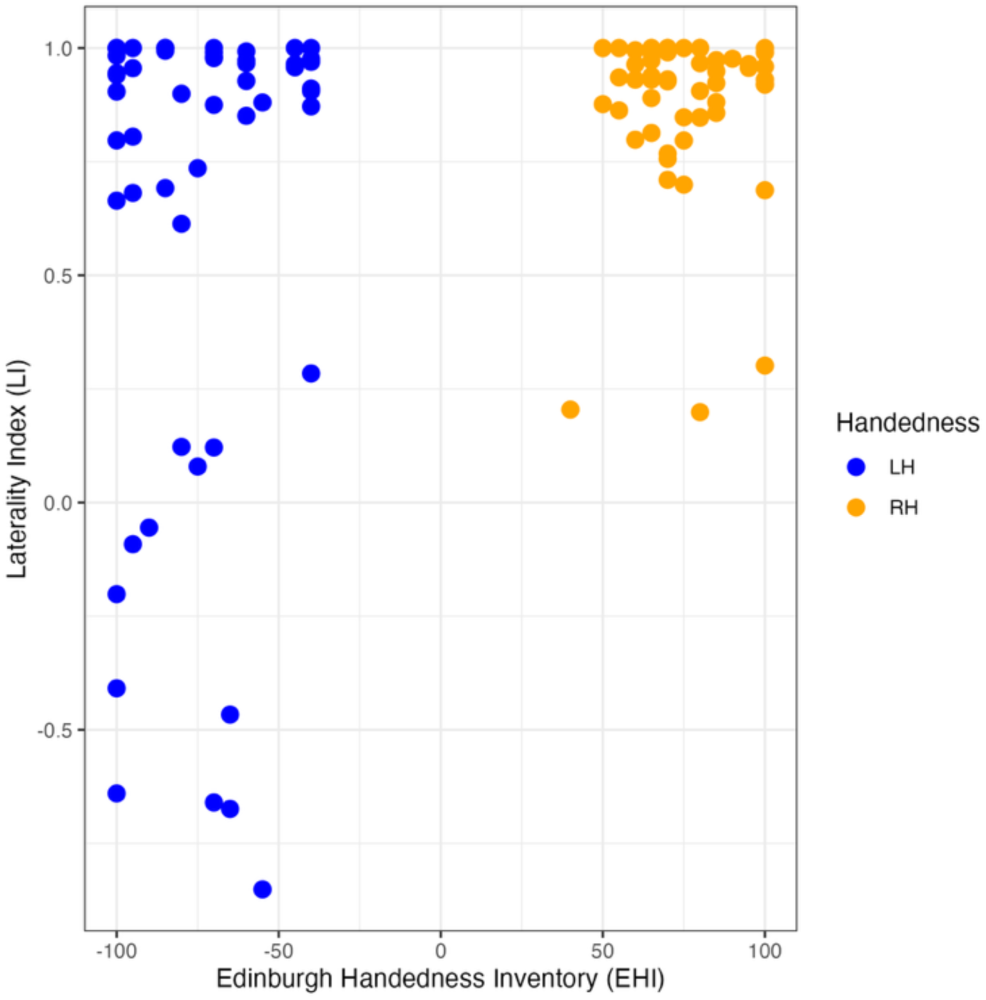
Language laterality indices (LI_lang_) and Edinburgh Handedness Inventory (EHI) scores for left- and right-handed participants. LH = left-handed; RH = right-handed

#### Memory-Related Activation

We contrasted activation for subsequent hits with subsequent misses in the word and face encoding conditions for LH and RH groups separately. For face encoding, we observed widespread activation for subsequent hits in ventral visual streams bilaterally as well as in medial temporal lobe, ventromedial prefrontal cortex, and right inferior frontal sulcus for both RH and LH groups (Figure 3A; see Supplementary Table for complete coordinates and cluster-level significance). For subsequent misses, there was widespread activation in the default mode network (DMN) for both groups, consistent with previous studies demonstrating subsequent memory effects (Shrager et al., 2008; Uncapher & Wagner, 2009). Critically, there were no significant activation differences when comparing groups directly.

**Figure 3.**
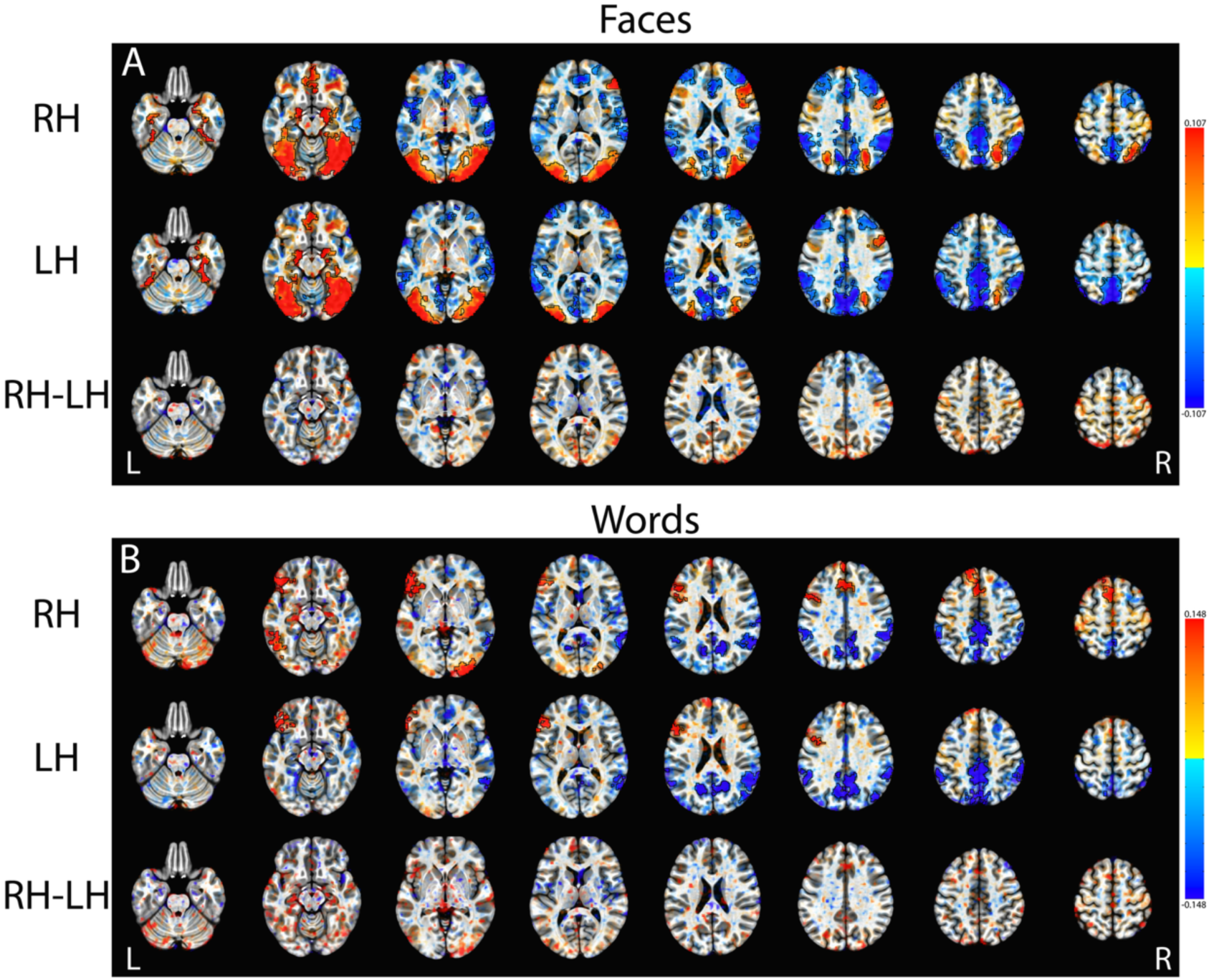
Subsequent memory contrast (subsequent hits > subsequent misses) for face stimuli (A) and word stimuli (B) in LH and RH groups. Contrasts were thresholded at voxel-wise p<0.01 (corrected p<0.05). Significant clusters are signified by black outlines. Note that there were no significant clusters in the RH-LH contrast. Left is on the left.

Subsequent memory effects for words were not as pronounced as those for faces (Figure 3B), likely due to near-ceiling memory performance for word recognition with very low trial counts for subsequent misses (mean = 8.5 miss trials across participants). However, consistent with subsequent memory effects for faces, we observed greater activation in DMN regions including precuneus and left and right angular gyrus for subsequently forgotten stimuli in both LH and RH groups. In contrast to the face contrast, for words we observed greater activation for subsequent hits in the inferior frontal gyrus on the left in both groups. Additionally, we observed significant activation for subsequent hits in the RH group in left inferior frontal sulcus, bilateral dorsomedial prefrontal cortex (DMPFC), and right lateral occipital lobe. Again, when comparing groups directly we observed no significant activation differences.

For the retrieval data, we again performed memory contrasts (hits vs. CRs) for faces and words separately. For faces, we observed widespread memory retrieval activations for both LH and RH groups (Figure 4A), with significant activation for hits in bilateral visual regions (including occipital, parietal, and temporal visual regions), frontal regions (bilateral inferior frontal sulcus, DMPFC, left frontopolar), and bilateral caudate nucleus. There were significant activations for CRs in bilateral VMPFC, right DLPFC, right middle temporal gyrus, and right primary visual cortex for the LH group only. Again, direct comparison of LH and RH groups yielded no significant activation differences. For word retrieval, we observed widespread cortical and subcortical activation for hits in both LH and RH groups (Figure 4B) and no significant activation differences between groups when directly contrasting LH and RH groups.

**Figure 4.**
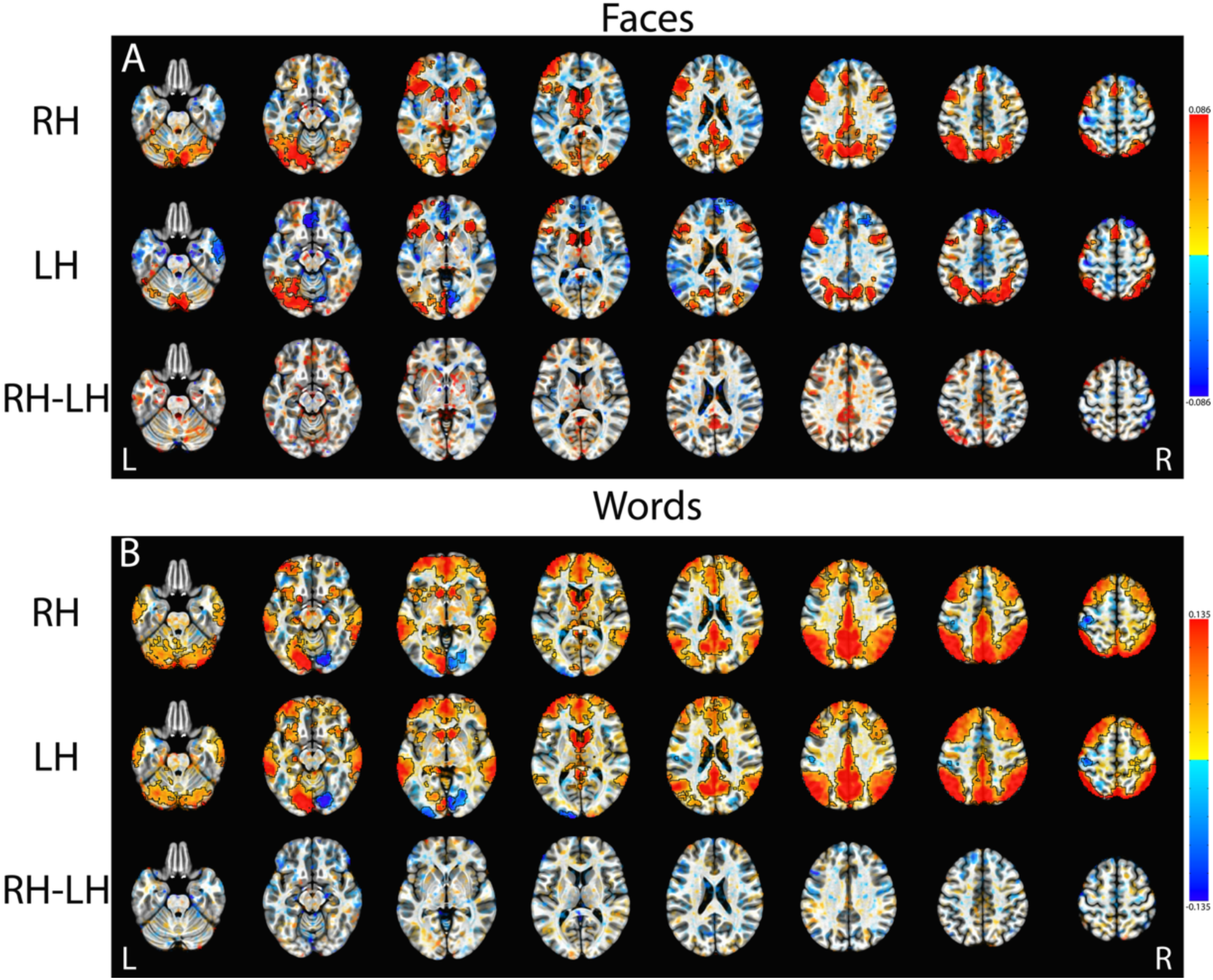
Significant clusters of activation in the contrast of hits > CRs for faces (A) and words (B) in the LH and RH groups. Contrasts were thresholded at voxel-wise p<0.01 (corrected p<0.05). Significant clusters are signified by black outlines. Note that there were no significant clusters in the RH-LH contrast. Left is on the left.

Given that we did not observe group differences when comparing the full LH and RH samples, we next asked whether we would observe differences in activation if we compared groups based on LI_lang_. Accordingly, we repeated the above analyses for comparing activation for participants with negative LI_lang_ scores (n=9) against all participants with positive LI_lang_ scores.

We observed activation differences for the word retrieval condition in the right DMPFC, right frontal pole, right inferior parietal lobule, and right lateral temporal lobe (Figure 5; Table 2). In each case, the right-lateralized group had greater activation differences than the left-lateralized group for word retrieval. We did not observe any group differences in any other task conditions.

**Figure 5.**
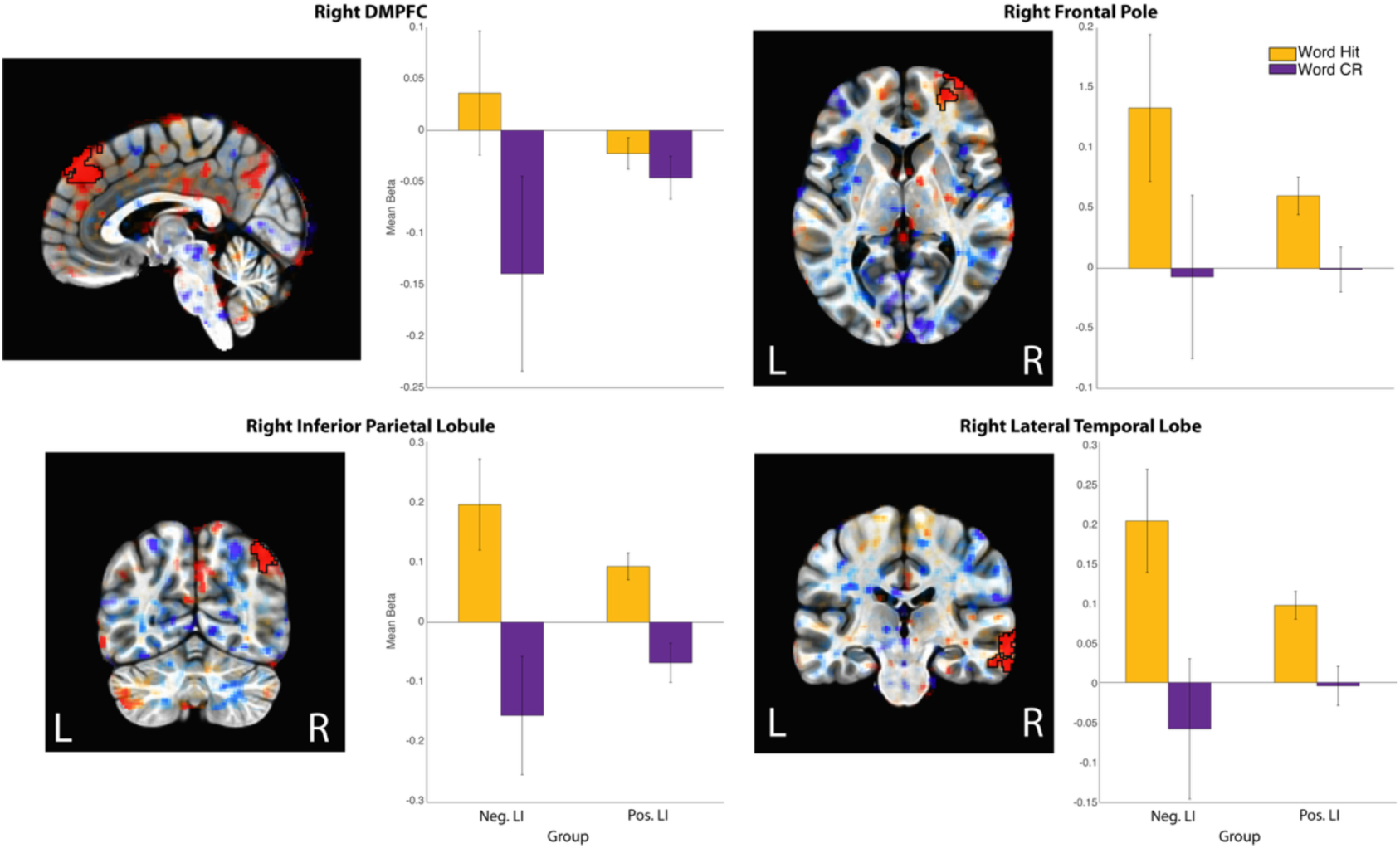
Significant clusters of activation in the group contrast between negative and positive LI_lang_ for the word retrieval task condition. Significant clusters were observed in the right DMPFC, right frontal pole, right inferior parietal lobule, and right lateral temporal lobe. Contrasts were thresholded at voxel-wise p<0.01 (corrected p<0.05). Left is on the left.

**Table 2.**
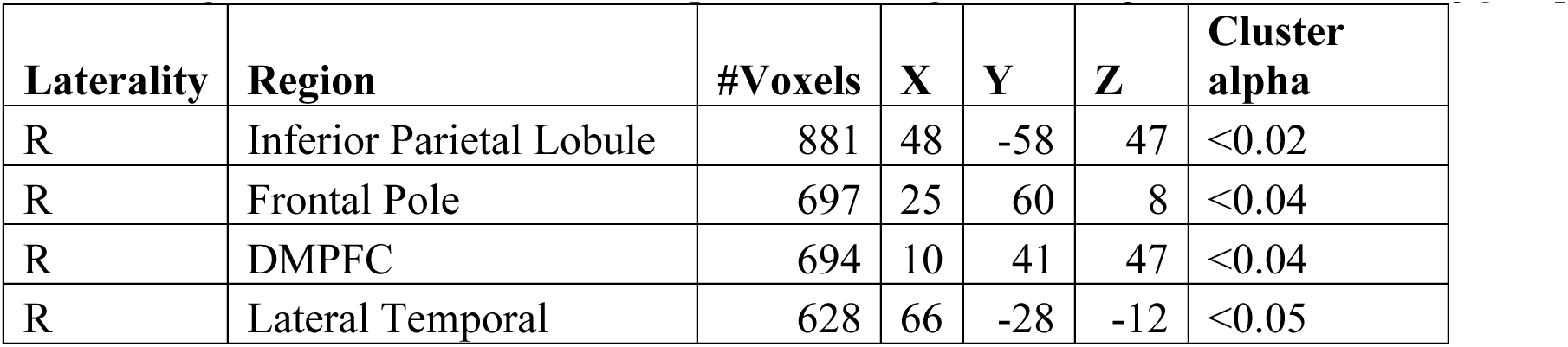
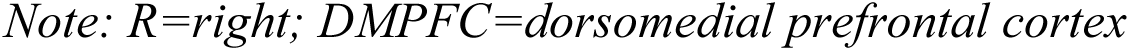
Significant clusters in the comparison of Negative-LI_lang_ and Positive-LI_lang_ groups.

| Laterality | Region | #Voxels | X | Y | Z | Cluster alpha |
| --- | --- | --- | --- | --- | --- | --- |
| R | Inferior Parietal Lobule | 881 | 48 | -58 | 47 | <0.02 |
| R | Frontal Pole | 697 | 25 | 60 | 8 | <0.04 |
| R | DMPFC | 694 | 10 | 41 | 47 | <0.04 |
| R | Lateral Temporal | 628 | 66 | -28 | -12 | <0.05 |

*Note: R=right; DMPFC=dorsomedial prefrontal cortex*

#### MTL Anatomical ROI Analysis

To investigate memory-related activation in the MTL, we examined average activation within hippocampus, entorhinal cortex, perirhinal cortex, and parahippocampal cortex in left and right hemisphere for both groups. Mixed ANOVAs with a between factor for group, and within factors for ROI and hemisphere were conducted for each memory task. In the face encoding task, there were significant main effects of hemisphere (F(1,107) = 9.80, p = 0.002). The main effect of ROI was significant in each task (p’s < 0.05). There were no significant main effects of group or significant interactions involving group. See Supplementary Table 2 for full ANOVA results.

#### Monte Carlo Simulations

Our final hypothesis was that including LH participants would increase the variability of results compared to all-RH samples. To test this, we performed Monte Carlo simulations where we randomly selected subsets of participants with sample sizes between 10 and 50 and included between 0 and 9 LH participants. Figure 6 presents the DSC values for each sample size and task condition when the proportion of LH participants (k) is approximately 10% of the sample (Gilbert & Wysocki, 1992). For DSC tables for all combinations and corresponding binomial probabilities, please see OSF (https://osf.io/f568s/overview?view_only=4f53cd74ad5646e68272ee9ff9e6b4a4). DSC values range from 0 to 1, where scores ≥ 0.7 are considered “good” and scores ∼0.8-0.9 are standard for medical image registration (Brock et al., 2017). In our Monte Carlo simulations, DCS scores were generally high, even when including up to 9 LH participants, especially when sample sizes were n ≥ 30. The word encoding condition had the lowest DSC scores generally, however, when considering only those combinations of sample size and number of LH participants that are statistically likely (e.g., those where the probability of selecting (k) LH participants in sample size (n) is ≥ 0.05), then DSC values are above approximately 0.6 and are as high as 0.9 with the largest sample sizes.

**Figure 6.**
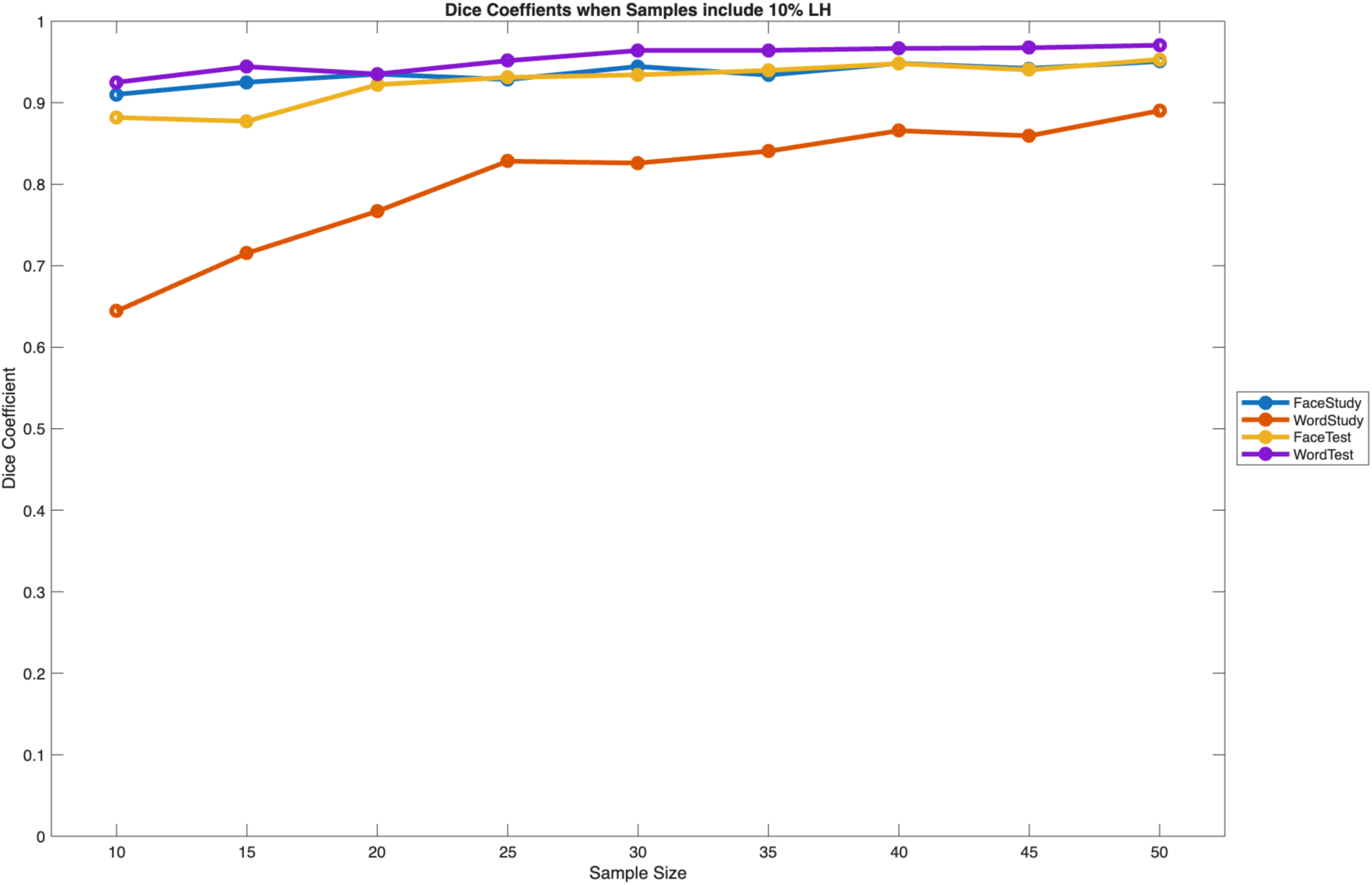
Dice similarity coefficients (DSC) between the activation maps when samples include all RH participants and when samples include approximately 10% LH participants. Smaller sample sizes had lower DSC values generally. All DSC values are very good when sample sizes are larger than ∼30.

We examined the effect of LI_lang_ on activation overlap in the Monte Carlo simulations for the largest sample sizes (n=50), comparing the all-RH samples (k=0) with samples including 10% LH (k=5) for all task conditions. Across all four memory tasks (face/word encoding and retrieval), pairwise similarity between Monte Carlo draws was driven predominantly by participant overlap between draws (Jaccard index; B = 0.403, SE = 0.002, t = 185.62). Once overlap was accounted for, there was no residual effect of inclusion of LH participants (k00 vs. k05; B = 0.0004, SE = 0.0016, t = 0.24) and no effect of the degree of LI_lang_ difference between specific draws (B = 0.007, SE = 0.075, t = 0.09), nor an interaction between the two (B = 0.004, SE = 0.077, t = 0.06). This indicates that the reduced map similarity observed when left-handed participants are included in the sampling pool is fully attributable to reduced subject overlap between draws (mean Jaccard index: k00 = 0.83, k05 = 0.60) rather than to any systematic effect of LI_lang_ on the underlying activation maps.

## Discussion

The overarching aim of this study was to examine whether there was a basis for exclusion of LH participants from fMRI memory research by determining whether there was a significant difference in brain activation between LH and RH participants when performing memory tasks. At a behavioral level, we saw no difference in performance between LH and RH groups for memory for words or for faces. Similarly, our whole-brain analyses revealed no effects of handedness on memory encoding or retrieval activation for either faces or words. The only group differences we observed were when we specifically compared participants who had atypical (i.e., right hemisphere) language lateralization with those who did not. In that case only word retrieval was affected with greater activation in right hemisphere clusters for the right-lateralized group but no increased activations evident for the left-lateralized group. Even when adding progressively more LH participants to our samples, we observed robust activation maps, especially with large sample sizes, and even when the samples comprised majority LH participants. For fMRI studies of memory processes, we suggest that these results support including larger sample sizes with representative numbers of left-handed participants (Bailey et al., 2020).

### Lateralized Memory Effects

Consistent with models of human memory that incorporate lateralization of effects (e.g., HERA; Nyberg et al., 1996), we observed lateralized effects during encoding and retrieval of faces and words in both RH and LH groups. One consistent finding was increased activation in the right inferior frontal sulcus for subsequent hits for faces (compared to subsequent misses). For both LH and RH groups there was more activation in this condition on the right than on the left, with left hemisphere activation failing to reach significance following corrections for multiple comparisons. Previous studies have demonstrated subsequent memory effects for faces in similar regions of frontal lobes (Keightley et al., 2011), with some also showing right-lateralized effects (Sergerie et al., 2005). Other studies have shown activation in similar regions for face processing more generally; for example in a probabilistic map of face processing in 1,110 adolescents (Tahmasebi et al., 2012) and face processing in parents of ASD children (Yucel et al., 2015). For word encoding, we observed left-lateralized subsequent memory effects in both LH and RH groups, consistent with meta-analyses of subsequent memory in fMRI (Kim, 2011).

At retrieval for face stimuli, we again observed inferior frontal regions were activated for hits compared to correct rejections. While there were significant clusters in both hemispheres, the left hemisphere cluster had a larger spatial extent in both LH and RH groups. Word retrieval similarly had larger extent on the left than on the right in both groups.

We also noted strong lateralized effects in both word and face retrieval conditions in the primary visual cortex (especially evident in Figure 4B). We interpret this result as a confound of our experimental design, as the response options “old” and “new” were displayed on the lower left and lower right, respectively, of the screen during all test trials. If, for example, participants looked at the lower left of the screen when selecting the “old” response for target stimuli, the upper right of their visual field would receive more stimulation resulting in more activity in visual neurons in the left hemisphere which have corresponding receptive fields. Future studies may wish to either include eye tracking measures or display response options in the central part of the display to minimize this confound.

### Variability due to including LH participants

The conventional wisdom is that including LH participants in fMRI studies adds variability to the data and increases the potential for false-negative results. In a sense, our results support this position; when sample sizes in our Monte Carlo simulations were small (e.g., n=10), the inclusion of just a few LH participants resulted in much lower overlap with the all-RH sample activation maps. However, when sample sizes were more robust (e.g., n ≥ 30), Dice similarity scores were greater than 0.8 for all contrasts. This result is consistent with the observation that small sample sizes are more prone to spurious results (Button et al., 2013) and further supports the argument for larger sample sizes in fMRI research. In a recent systematic review (Monsour et al., submitted), the median sample size in the fMRI memory literature was n=32. Consequently, our results indicate that the inclusion of LH participants would not have a strong effect on the results of most memory fMRI studies.

The condition where we saw the greatest effect of including LH participants was word encoding. We urge caution in interpreting this result, as recognition memory performance for words in our study was near ceiling, meaning that there were very few trials in the “subsequent miss” category. Accordingly, future studies may wish to modify the experimental design to increase the number of subsequent misses (e.g., by increasing list lengths or using longer delays) to determine whether our results are due to lower inherent contrast between task conditions or due to handedness-related laterality effects. In any case, we note as above that with larger sample sizes results in this condition were still robust.

### Handedness and fMRI Activation

Our results contrast with those of Cuzzocreo et al. (2009), who found greater activation in LH than RH participants for both encoding and retrieval of paired associate words in the left MTL. Several differences between task design and groups may account for this discrepancy.

First, our task employed single item encoding and retrieval, which may place lower demands on the MTL/hippocampus. We also sampled young adults whereas the previous study examined memory processes in older adults. We note that, consistent with our results, there was not an absence of activation in the LH group, but rather additional activation relative to the RH group for verbal memory tasks.

Other studies have observed activation differences associated with handedness in both resting-state and task-based fMRI designs. For example, Tejavibulya and colleagues (2025) examined resting state functional connectivity in large cohorts included in the Human Connectome Project (HCP) and found significant differences between RH and LH participants in both the HCP-Developmental cohort (n = 465; ages = 5-21 years) and in the HCP-Aging cohort (n = 368; ages = 36-100 years), though, notably, the exact locations of differences between groups differed between the age ranges and between items used to assess handedness.

Badzakova-Trajkov and colleagues (2010) scanned 48 LH and 107 RH participants while they completed word-generation, spatial attention, and face processing tasks. Laterality indices were calculated for each processing domain. Consistent with previous studies (Isaacs et al., 2006; Knecht et al., 2000; Pujol et al., 1999; Szaflarski et al., 2002), language was left-lateralized in 95% of RH participants but only 81% of LH participants. Face processing also demonstrated handedness-related lateralization, with 94% and 73% of RH and LH participants showing right-lateralized activation, respectively (Bourne, 2008). Interestingly, spatial attention did not demonstrate a relationship between handedness and lateralization, with 79% of both groups being right lateralized for this task. This is consistent with a recent large-scale behavioral study that showed no differences in spatial navigation ability between LH and RH participants (Fernandez-Velasco et al., 2023). Interestingly, a recent fMRI study found that LH participants with left-lateralized language processing exhibited less lateralization of function (i.e., lower LI values on average) compared to RH participants for visual processing of bodies, faces, and scenes. Finally, a recent study suggests that in participants with right LI_lang_ also exhibited atypical lateralization for face processing, tool praxis, spatial attention, and emotion prosidy (Gerrits et al., 2020). In our contrasts based on LI_lang_, we did not observe significant activation differences in face recognition. Future studies may wish to pre-select participants with right LIlang as in Gerrits et al. (2020) to ensure adequate statistical power to investigate this question further.

### Limitations

There are additional limitations of our study that should be considered. First, all participants, regardless of handedness, were required to respond to all behavioral tasks with their right hand. There is evidence that fine motor control, specifically for sequential finger movements, is differentially lateralized for RH and LH individuals (Solodkin et al., 2001).

Because of the lack of research in this area, it is unclear whether having LH individuals respond with their non-dominant hand resulted in activation differences compared to the RH group. We note that there were no significant activation differences between groups in any motor regions. Further, there were no reaction time differences between groups for any of the task conditions, indicating that response selection with the non-dominant hand did not add significant processing time for the LH group. Second, the observed effects may be selective to words and faces and may not generalize to other stimulus types. Further, we did not calculate a separate face laterality index, which may have been necessary to observe group differences in face encoding or retrieval activation. However, Badzakova-Trajkov and colleagues (2010) demonstrated strong intercorrelations between handedness and laterality indices for both language and faces, indicating that any group differences in lateralization for face processing may have been captured by our LI_lang_. Third, we did not include ambidextrous individuals in our sample as we selected for those with clear handedness preferences. Prior research has shown language lateralization to be correlated with self-reported handedness (Isaacs et al., 2006; Szaflarski et al., 2002). Consequently, we chose to set cutoff scores for inclusion in LH and RH groups to give the greatest chance of observing group differences. Given the similarity of activation for LH and RH groups, we predict individuals without clear hand preferences for manual tasks would also show similar activations. Future studies may wish to test this prediction.

Behavioral performance for the word retrieval task was much better than for the face retrieval task and was near ceiling. This resulted in very few subsequent miss trials in the word encoding condition, potentially affecting the strength of that fMRI contrast. Our experimental design attempted to match the two tasks in terms of list length and delay between encoding and retrieval. Prior research has also observed better performance for words than faces (for review, see Robotham & Starrfelt, 2017). In our own research, we have observed better memory for objects than for faces in similar recognition memory tasks (Kirwan & Stark, 2007). We attribute these behavioral performance differences to both differences in cognitive processes involved in the initial processing of verbal and non-verbal materials as well as to differences in between-item interference for faces (which are all the same semantic category) compared to words. Future studies may wish to match behavioral performance between stimulus types through manipulations such as changing retention delays or list lengths.

Finally, while our sample size for both LH and RH groups was larger than typical for task-based fMRI studies (Yeung, 2018), we still may have lacked statistical power to detect differences between groups in either behavioral or fMRI measures. Indeed, we note that there were several significant clusters of activation in one group that did not reach significance in the other group (Figures 3 & 4). It is unclear whether these differences reflect variation due to the particular sample or whether they reflect group differences with smaller effect sizes. Larger study samples will be needed to rule out this possibility. We note that this is not a concern for the Monte Carlo simulations, which explicitly addressed sampling variability.

### Recommendations and Conclusions

We asked whether there were systematic differences in memory-related brain activation between left- and right-handed participants. At the whole brain level, we did not observe any activation differences when comparing participants based on self-reported handedness. We only observed activation differences when splitting groups according to language lateralization, and only in word retrieval tasks where we observed increased right-lateralized activation in participants with atypical (right) lateralization. Importantly, we did not observe any corresponding decreases in activation in the left hemisphere for these participants. When considering whether the addition of LH participants added variance to the data that diluted effects, we found robust overlap of activation maps, particularly for larger sample sizes.

If study samples are randomly selected from the population, then one can expect about 10% LH participants, of whom 75% will have typical lateralization for language and face processing (Badzakova-Trajkov et al., 2010). In other words, 10 out of 100 people are expected to be LH, and of those, on average only 2.5 will have atypical lateralization. In that same group of 100, 90 will be RH and of those 4.5 (or 5%) are expected to have atypical lateralization. Thus, in a large random sample that includes both RH and LH participants, atypical lateralization would be more frequently encountered in RH than in LH participants. As left-handedness is a common variant in the normal population, including LH participants in research samples is important if our results are to generalize to the larger population. If researchers choose to include LH participants, we recommend reporting handedness (either self-reported or measured with instruments such as the EHI) along with other demographic information. We echo previous authors (Bailey et al., 2020; Willems et al., 2014) in calling for cognitive neuroscience researchers to include left-handed participants in their research samples.

## Data availability statement

MRI data are available here: https://openneuro.org/datasets/ds004589. Code and supplemental methods are available here: https://osf.io/f568s/

## Ethics Approval

The research described here was approved by the Brigham Young University Institutional Review Board.

## Declaration of AI use

The authors used AI-assisted technologies in scripting data analyses. No AI-assisted technologies were used in writing the manuscript.

## Author contributions

LG and CBK conceived the research, designed and implemented the paradigm, and collected the data. LG, IH, and CBK analyzed the data and prepared the manuscript for publication.

